# Evidence of fitness costs associated with Ace-*1^R^*-mediated resistance to pirimiphos-methyl in *Anopheles gambiae* sensu lato following the withdrawal of indoor residual spraying in Burkina Faso

**DOI:** 10.64898/2026.08.07.743444

**Authors:** Delphine O. Karama, Aristide S. Hien, Dieudonné D. Soma, Kelly L. Ngaffo, Samina Maiga, Didier P. Alexandre Kaboré, Rabila Bamogo, Benson G. Meda, Moussa Namountougou, Roch K. Dabiré

**Affiliations:** Institut de Recherche en Sciences de la Santé, Direction Régionale de l’Ouest; BP 545 Bobo-Dioulasso, Burkina Faso; Université Nazi Boni, Bobo-Dioulasso, Burkina Faso

**Keywords:** *Ace-1*, *Anopheles gambiae* s.l., biological cost, pirimiphos-methyl, indoor residual spraying, insecticide resistance, Burkina Faso

## Abstract

**Introduction:** Changes in vector control strategies alter the selection pressures exerted on natural populations of *Anopheles gambiae* s.l. and may influence the dynamics of insecticide resistance mechanisms. However, the consequences of discontinuing indoor residual spraying (IRS) campaigns on the evolution of *Ace-1*-mediated resistance remain poorly documented under natural conditions. This study aimed to assess the spatiotemporal evolution of resistance to pirimiphos-methyl following the cessation of IRS and to investigate evidence consistent with the existence of a biological cost associated with *Ace-1*-mediated resistance.

**Methods:** Natural populations of *An. gambiae* s.l. were collected between 2017 and 2023 in three districts in Burkina Faso that had undergone IRS campaigns (Kampti, Solenzo, and Kongoussi). Susceptibility tests with pirimiphos-methyl (0.25%) were conducted out following WHO protocols, and a subsample of exposed mosquitoes was genotyped to detect the *Ace-1* G119S mutation. Spatiotemporal trends in mortality, allele frequencies and genotypes were analyzed according to the pre-IRS, IRS, and post-IRS periods. The association between the *Ace-1* genotype and survival following exposure to pirimiphos-methyl was assessed using logistic regression, while the concordance between phenotypic and molecular indicators of resistance was examined using Spearman’s correlation.

**Results:** The susceptibility of *An. gambiae* s.l. populations to pirimiphos-methyl was gradually restored after the discontinuation of IRS at all sites. At the same time, the frequencies of the *Ace-1* 119S resistance allele declined sharply, particularly in Kampti and Solenzo, while they remained low in Kongoussi throughout the study period. Mosquitoes carrying resistant genotypes had a significantly higher probability of survival after exposure to pirimiphos-methyl than susceptible homozygotes, with resistant homozygotes (RR) exhibiting the greatest survival advantage (OR = 27.62; 95% CI: 6.91-110.45; p < 0.001). A significant negative correlation was observed between the frequency of the *Ace-1* 119S allele and phenotypic mortality (ρ = −0.48; p = 0.033), indicating a concordance between the two indicators of resistance. The progressive decline in allele frequencies, the decreasing prevalence of resistant genotypes, and the concomitant restoration of susceptibility are field observations consistent with the existence of biological costs associated with *Ace-1*-mediated resistance.

**Conclusion:** This study provides field evidence consistent with the existence of biological costs associated with *Ace-1*-mediated resistance in natural populations of *An. gambiae* s.l. These results underscore the value of an adaptive resistance management strategy based on alternating selection pressures and could guide future vector control strategies, particularly if indoor residual spraying campaigns or other interventions relying on organophosphates were reintroduced.

## INTRODUCTION

Organophosphate-based indoor residual spraying (IRS), particularly with pirimiphos-methyl, has become a key component of insecticide resistance management strategies across sub-Saharan Africa following the widespread emergence of pyrethroid resistance in malaria vectors [1, 2]. However, sustained exposure to organophosphates exerts strong selective pressure on mosquito populations, favoring the spread of resistance mechanisms that compromise the efficacy of these interventions [3, 4]. Among these mechanisms, the G119S substitution in the acetylcholinesterase gene (*Ace-1*) is one of the most important target-site mutations conferring resistance to both organophosphate and carbamate insecticides [5]. This mutation reduces the sensitivity of acetylcholinesterase to insecticide inhibition, thereby enhancing mosquito survival following exposure to compounds such as pirimiphos-methyl [6]. The *Ace-1* 119S allele has been reported in several West African populations of *Anopheles gambiae* s.l. and is strongly associated with phenotypic resistance to organophosphates, raising concerns regarding the long-term sustainability of IRS programmes relying on these insecticides [7, 8].

Although insecticide resistance alleles provide a selective advantage in treated environments, their persistence in the absence of insecticide pressure is often constrained by fitness costs [8, 9]. Experimental studies have demonstrated that the *Ace-1* 119S mutation can impair the catalytic efficiency of acetylcholinesterase, leading to reduced fitness-related traits such as survival, development, reproductive performance, and overall competitiveness compared with susceptible mosquitoes [6, 10]. Consequently, when insecticide pressure is removed, susceptible alleles may progressively recover in frequency through natural selection [8, 6]. In *Anopheles gambiae*, evidence from laboratory and field studies suggests that the fitness burden associated with resistant *Ace-1* genotypes can drive a decline in resistance frequencies over time, although this process may be moderated by gene duplication events that partially compensate for the deleterious effects of the mutation [3, 6, 10, 11]. Understanding how *Ace-1* frequencies change following the withdrawal of IRS therefore provides a valuable opportunity to assess the evolutionary dynamics of resistance and to infer the magnitude of associated fitness costs under natural field conditions [3, 4, 12].

This study aimed to determine whether the withdrawal of pirimiphos-methyl-based indoor residual spraying (IRS) was associated with a decline in Ace-1-mediated resistance in *Anopheles gambiae* s.l. populations in Burkina Faso and to assess whether observed changes were consistent with fitness costs associated with the Ace-1 resistance allele. Specifically, we investigated temporal changes in phenotypic susceptibility to pirimiphos-methyl, the frequency of the *Ace-1* 119S resistance allele, and genotype–phenotype associations before, during and after IRS implementation.

## MATERIAL AND METHODS

### Study area

The study was conducted across Burkina Faso, a landlocked West African country characterized by three major eco-climatic zones that differ markedly in rainfall patterns, vegetation cover, agricultural activities and malaria transmission intensity. From north to south, these zones comprise the Sahelian zone, the Sudan–Sahelian zone and the Sudanian zone. The Sahelian zone occupies the northern part of the country and is characterized by low annual rainfall (<600 mm), sparse vegetation and a short transmission season. The Sudan–Sahelian zone, located in the central region, represents a transitional ecological belt with intermediate rainfall (600–900 mm) and moderate agricultural activity. The Sudanian zone in the south receives the highest annual rainfall (>900 mm), supports dense vegetation and extensive agricultural production, and generally experiences longer malaria transmission seasons.

A network of insecticide resistance monitoring sites was established across the three eco-climatic zones to provide nationwide surveillance of malaria vector susceptibility to insecticides. A total of 27 sentinel sites were distributed throughout the country, encompassing diverse ecological settings and malaria transmission profiles. In the Sahelian zone, monitoring sites included Dori, Ouahigouya, Séguénéga, Kaya and Kongoussi. Within the Sudan–Sahelian zone, surveillance was conducted in Nouna, Dédougou, Ouagadougou, Koudougou, Boromo, Houndé, Manga, Tenkodogo, Fada N’Gourma and several surrounding localities. In the Sudanian zone, monitoring activities were implemented in Vallée du Kou, Bobo-Dioulasso, Orodara, Banfora, Tiéfora, Karangasso-Vigué, Soumousso, Diébougou, Gaoua, Kampti, Dano and Mangodara. Three sites were selected for the implementation and evaluation of pirimiphos-methyl-based indoor residual spraying (IRS) interventions. These sites were strategically located in areas with high malaria transmission and operational relevance for vector control programmes. IRS implementation sites included Kongoussi in the northern Sahelian zone, Solenzo in the western Sudan–Sahelian zone and Kampti in the southern Sudanian zone. The distribution of these IRS sites across the three major eco-climatic zones provided an opportunity to evaluate the impact of insecticide-based interventions under contrasting ecological conditions and varying levels of vector exposure to insecticide selection pressure (fig.1).

**Figure 1.**
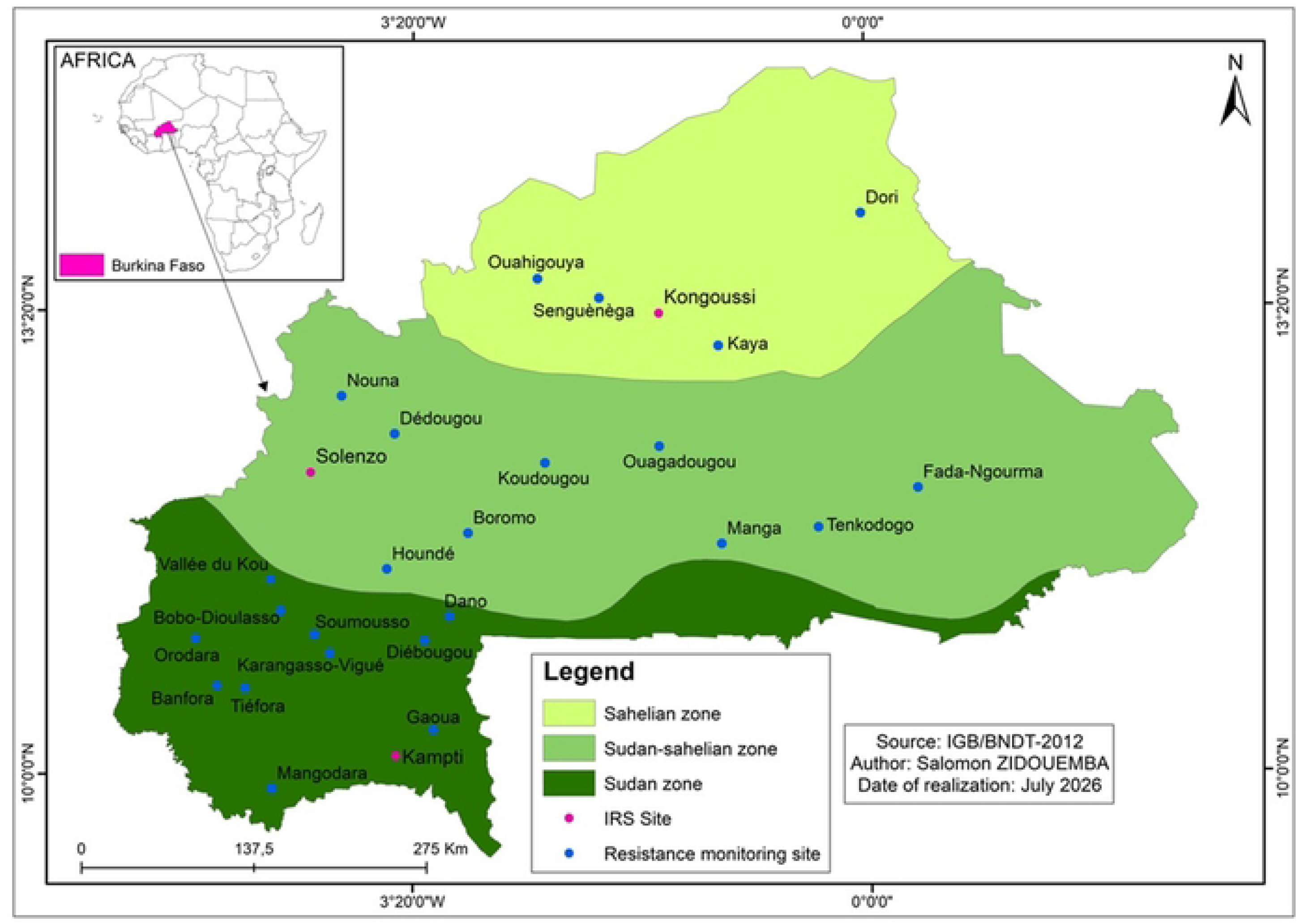
Geographic distribution of indoor residual spraying (IRS) implementation sites (pink dots) and insecticide resistance monitoring sites (blue dots) across Burkina Faso (2017-2023). The map illustrates the three major eco-climatic zones of the country: the Sahelian zone (light green), the Sudan–Sahelian zone (medium green), and the Sudanian zone (dark green). IRS implementation sites were located in Kongoussi, Solenzo, and Kampti, while insecticide resistance monitoring was conducted through a nationwide network of sentinel sites distributed across the three ecological zones.

### IRS implementation history

Table 1 summarizes the history of indoor residual spraying (IRS) in the three study districts of Kongoussi, Solenzo and Kampti, where IRS campaigns based on the organophosphate insecticide pirimiphos-methyl (Actellic® 300CS) were deployed as part of the national malaria vector control programme. The staggered implementation and subsequent withdrawal of IRS across these sites provided a valuable natural experimental framework for assessing the long-term impact of insecticide selection pressure on vector populations and the evolution of insecticide resistance.

**Table 1.**
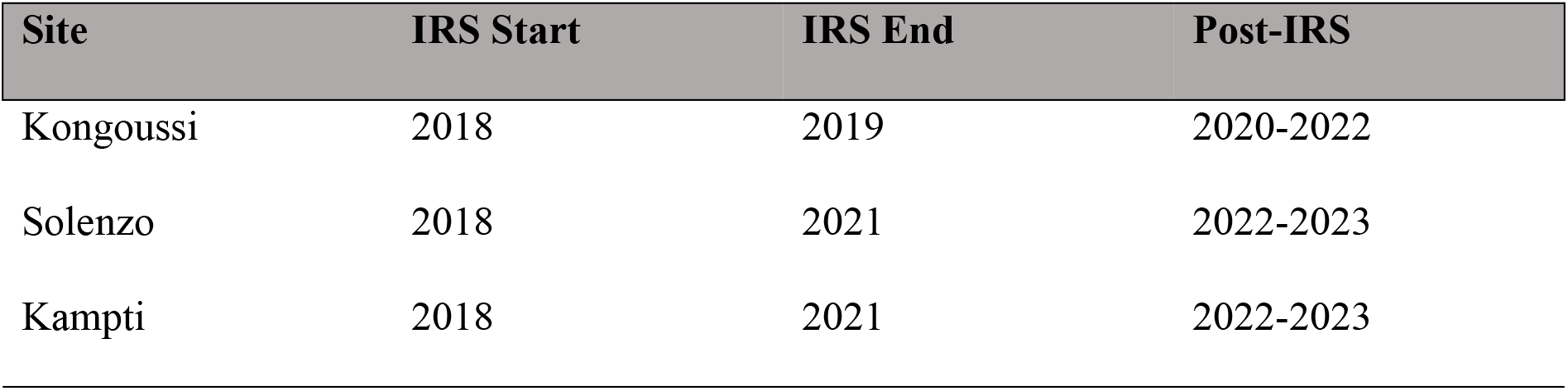
History of Indoor Residual Spraying in the three study sites districts.

In Kongoussi, IRS with Actellic® 300CS was implemented for two consecutive years (2018-2019), after which spraying activities were discontinued. This resulted in a relatively long post-IRS period spanning three years (2020-2022), offering an opportunity to investigate the persistence of resistance mechanisms in the absence of continued organophosphate exposure. Such an extended withdrawal period is particularly suitable for evaluating whether resistance alleles, including the *Ace-1* G119S mutation associated with organophosphate resistance, decline over time due to fitness costs when insecticide selection pressure is relaxed. In Solenzo and Kampti, Actellic® 300CS was applied at least once, followed by other formulations between 2018 and 2021, corresponding to four consecutive spraying campaigns. The longer duration of indoor residual spraying (IRS) in these two districts likely exerted stronger and more sustained selective pressure on local populations of *An. gambiae* s.l., potentially favoring the accumulation and maintenance of alleles associated with resistance (table.1). Consequently, the shorter post-IRS period (2022-2023) in Solenzo and Kampti provides an important contrast with Kongoussi for examining the temporal dynamics of resistance decline following the cessation of IRS. The variation in IRS duration among the three sites creates a gradient of historical insecticide exposure, ranging from moderate exposure in Kongoussi (two years of IRS) to more intense and prolonged exposure in Solenzo and Kampti (four years of IRS). Such differences are expected to influence both phenotypic susceptibility and the frequency of resistance-conferring alleles within mosquito populations. Therefore, comparing vector populations across these sites provides a robust framework for assessing the relationship between historical IRS intensity, changes in *Ace*-*1*^R^ allele frequencies, and the potential fitness costs associated with pirimiphos-methyl resistance.

Overall, the IRS implementation history indicates that all three study sites were exposed to at least one annual round of Actellic® 300CS IRS, ensuring substantial organophosphate selection pressure on local malaria vectors. The subsequent discontinuation of spraying created a unique opportunity to evaluate whether the removal of pirimiphos-methyl-based IRS leads to a reduction in Ace-1-mediated resistance, thereby generating field evidence for fitness costs associated with organophosphate resistance in *Anopheles gambiae* s.l. populations in Burkina Faso.

### Study design

This study was designed as a longitudinal natural evolutionary experiment aimed at assessing whether the withdrawal of pirimiphos-methyl-based indoor residual spraying (IRS) was associated with changes in phenotypic susceptibility and *Ace-1*-mediated resistance in natural populations of *An. gambiae* s.l. in Burkina Faso. Unlike laboratory selection studies, this approach exploited operational vector control interventions implemented as part of routine national malaria control activities, which allowed resistance dynamics to be followed under real-world conditions.

The study encompassed three distinct periods corresponding to different levels of insecticide selection pressure: (i) the pre-IRS period (2017), during which only routine insecticide resistance monitoring was conducted prior to the implementation of IRS; (ii) the IRS period (2018–2021, depending on the study site), during which annual rounds of indoor residual spraying with the organophosphate insecticide pirimiphos-methyl (Actellic® 300CS) were implemented alongside routine resistance monitoring; and (iii) the post-IRS period (2020-2022 in Kongoussi; 2022-2023 in Solenzo and Kampti), characterized by the cessation of IRS activities while resistance surveillance continued. In 2023, no collection could be carried out in Kongoussi due to insecurity, which made the site inaccessible.

The conceptual framework underpinning this design is based on evolutionary theory predicting that insecticide resistance alleles increase in frequency under sustained insecticide selection but may decline when selection pressure is removed if they impose fitness costs on their carriers. During the IRS phase, repeated exposure of mosquito populations to pirimiphos-methyl was expected to favor the survival and reproduction of individuals carrying the resistant *Ace-1* 119S allele, resulting in elevated frequencies of resistant genotypes and reduced phenotypic susceptibility. Following the withdrawal of IRS, however, the absence of organophosphate exposure was expected to relax selection pressure, thereby enabling susceptible genotypes to regain a competitive advantage if resistant mosquitoes experienced reduced biological fitness. To test this hypothesis, temporal trends in phenotypic mortality following exposure to pirimiphos-methyl and changes in the frequency of the *Ace-1* G119S mutation were monitored throughout the study period. Indirect evidence for fitness costs was inferred from a combination of evolutionary and epidemiological indicators, including: (i) progressive increases in mortality rates in WHO susceptibility bioassays during the post-IRS period; (ii) concomitant declines in the frequency of the resistant *Ace-1* allele; and (iii) changes in genotype–phenotype associations over time. Under a fitness-cost scenario, withdrawal of IRS would be expected to result in a gradual restoration of susceptibility, reflected by increasing phenotypic mortality and decreasing prevalence of resistant genotypes. Conversely, persistence of resistance despite the removal of insecticide pressure would suggest either negligible fitness costs or the existence of compensatory mechanisms, such as *Ace-1* gene duplication, capable of mitigating the deleterious effects associated with the resistant allele.

The inclusion of multiple study sites with contrasting IRS histories further strengthened the inference framework. Kongoussi, which experienced a longer post-IRS period, provided an opportunity to evaluate long-term evolutionary responses following the cessation of selection pressure, whereas Solenzo and Kampti enabled assessment of resistance dynamics after more prolonged IRS exposure.

### Mosquito collection

Immature stages of *Anopheles gambiae* sensu lato (*An. gambiae* s.l.) were collected annually from established insecticide resistance monitoring sites between July and November, corresponding to the peak malaria transmission season, from 2017 to 2023. Sampling was conducted as part of the national insecticide resistance surveillance program to ensure consistent monitoring of vector susceptibility trends before, during, and after the implementation of indoor residual spraying (IRS) interventions. Larval collections were performed in a wide range of natural and human-made aquatic habitats commonly exploited by *An. gambiae* s.l., including puddles, drainage channels, stream-bed pools, temporary rain-fed pools, swamps, brick-making pits, and irrigated rice fields. Standard dipping techniques were used to collect larvae from all available breeding sites within each sentinel locality. To minimize sampling bias and maximize representation of the local vector population, larvae from multiple breeding habitats were pooled at the site level prior to laboratory rearing. Collected larvae were transported to the insectary of the Institut de Recherche en Sciences de la Santé, Direction Régionale de l’Ouest (IRSS-DRO) Bobo-Dioulasso, where they were maintained under standardized environmental conditions (27 ± 2°C, 70 ± 10% relative humidity, and a 12:12 h light photoperiod). Larvae were reared to adulthood following established WHO procedures for mosquito husbandry. Emerging adults were provided with 10% sucrose solution ad libitum until use in bioassays. Adult mosquitoes were morphologically identified using standard taxonomic keys, and only female *An. gambiae* s.l. aged 2–5 days that had not previously blood-fed were selected for insecticide susceptibility testing.

### Phenotypic Susceptibility Assays to Pirimiphos-Methyl

Phenotypic susceptibility of *An. gambiae* s.l. populations to the organophosphate insecticide pirimiphos-methyl was evaluated annually from 2017 to 2023 using WHO tube bioassays performing according to standard procedures [13]. These bioassays were conducted using insecticide-impregnated papers containing the WHO diagnostic concentration of pirimiphos-methyl (0.25%), which is specifically designed to discriminate between susceptible and resistant phenotypes within mosquito populations. Control assays were performed simultaneously using papers treated only with paraffin oil. For each mosquito population, batches of approximately 100 non-blood-fed female mosquitoes (20-25 per tube) aged 2–5 days were exposed to pirimiphos-methyl-treated papers for 60 minutes, while a corresponding control group of 25 mosquitoes (for two tubes) was exposed to untreated papers. Following exposure, mosquitoes were transferred to holding tubes and maintained under standard insectary conditions with continuous access to a 10% s ucrose solution. Mortality was recorded 24 hours post-exposure in accordance with WHO guidelines [13]. Mosquitoes that died within the observation period were classified as phenotypically susceptible, whereas those surviving exposure were considered phenotypically resistant. Both dead and surviving mosquitoes were individually preserved in silica gel for subsequent molecular characterization of resistance markers, including the *Ace-1* G119S mutation. The preserving both surviving and dead specimens allowed direct genotype–phenotype association analyses.

### Molecular Characterization of Vector Species and *Ace-1^R^*-Mediated Resistance DNA Extraction and Species Identification

To investigate the molecular basis of temporal changes in pirimiphos-methyl susceptibility, genomic DNA was extracted from a subsample of 50 to 100 mosquitoes per site, comprising between 25 and 50 surviving mosquitoes (phenotypically resistant) and between 25 and 50 dead mosquitoes (phenotypically susceptible) following susceptibility testing. DNA was extracted from mosquito head-thorax specimens (preserved in silica gel) using the 2% cetyltrimethylammonium bromide (CTAB) protocol as previously described [14]. Because members of the *Anopheles gambiae* complex differ in their ecology, behavior, insecticide susceptibility profiles, and capacity to develop resistance, all specimens were identified to species level prior to resistance marker analysis. Species identification was performed using the SINE200 polymerase chain reaction (PCR) assay developed by Santolamazza et al. [15], which enables reliable discrimination between the principal malaria vectors within the *An. gambiae* complex. Species composition was determined throughout the study period in order to distinguish temporal changes in phenotypic resistance that reflected shifts in species distribution from those reflecting evolutionary change within vector populations.

### Detection of the *Ace-1* G119S Resistance Mutation

To evaluate the role of target-site resistance in the observed phenotypic responses to pirimiphos-methyl, mosquitoes were genotyped for the *Ace-1* G119S mutation, a well-established molecular marker associated with resistance to organophosphate and carbamate insecticides. Genotyping was performed using polymerase chain reaction–restriction fragment length polymorphism (PCR-RFLP) according to the protocol described by Weill et al. [16]. Amplifications were carried out in a final reaction volume of 25 μL containing 2.5 μL of 10× PCR buffer, 200 μM of each deoxynucleoside triphosphate (dNTP), 1 U/μL Taq DNA polymerase, 10 pmol of each primer (Ex3AGdir: 5′-GATCGTGGACACCGTGTTCG-3′ and Ex3AGrev: 5′-AGGATGGCCCGCTGGAACAG-3′) and 1-10ng of genomic DNA template. PCR amplification consisted of an initial denaturation at 94°C for 5 min, followed by 35 cycles of 94°C for 30 s, 54°C for 30 s, and 72°C for 30 s, with a final extension step at 72°C for 5 min. Amplified products were subsequently digested using 5 U of the restriction enzyme *AluI* and separated by electrophoresis on 2% agarose gels stained with ethidium bromide. DNA fragments were visualized under ultraviolet illumination, allowing discrimination among homozygous susceptible (*Ace-1SS*), heterozygous (*Ace-1RS*), and homozygous resistant (*Ace-1RR*) genotypes.

### Statistical analyses

All statistical analyses were performed using R software version 4.4.1 (R Foundation for Statistical Computing, Vienna, Austria). Statistical significance was assessed at the 5% level. Model assumptions were systematically evaluated using residual diagnostics and goodness-of-fit tests. Variations in the species composition of the *An. gambiae* s.l. complex were analyzed using multinomial logistic regression, with species as the response variable and the intervention period and site as fixed effects. Adjusted odds ratios (OR) and their 95% confidence intervals (95% CI) were estimated, and site × period interactions were assessed using likelihood ratio tests (LRT). For phenotypic sensitivity to pirimiphos-methyl, it was analyzed using generalized linear binomial models with a logit link, modeling the number of dead mosquitoes among those exposed according to intervention period and site. Differences in mortality between periods were assessed using ORs and 95% CI, with likelihood ratio tests. Temporal and spatial variations in the frequency of the *Ace-1* 119S resistance allele were studied using generalized linear binomial models, considering the number of resistance alleles relative to the total number of genotyped alleles. The effects of year, treatment period, species, and site were evaluated using models that included additive effects and interactions to examine the heterogeneity of temporal dynamics across sites. Estimated marginal means and pairwise comparisons were used to characterize variations in allele frequencies. Finally, the association between the *Ace-1* genotype and survival following exposure to pirimiphos-methyl was evaluated using logistic regression, by estimating the ORs and 95% CIs associated with the different genotypes. The correlation between the decrease in the frequency of the *Ace-1* 119S allele, the increase in phenotypic mortality and the gradual disappearance of resistant genotypes was considered field evidence consistent with the existence of biological costs associated with *Ace-1*-mediated resistance.

## RESULTS

### Species composition of *Anopheles gambiae* complex populations

A total of 1,098 specimens were identified by molecular assays between 2017 and 2023 across the three study sites. Of these, 298 originated from Kongoussi (Sahelian zone), 400 from Solenzo (Sudan-Sahelian) and 400 from Kampti (Sudanian zone). All three species of the *An. gambiae* complex were identified: *An. gambiae* s,s,, *An. coluzzii*, and *An. arabiensis* with predominating of *An. gambiae* s,s, throughout the study. Across period and sites, substantial variation in species composition was observed (fig.2). In Kongoussi, vector populations were initially dominated by *An. coluzzii* before and during IRS but a progressive shift toward codominance of this species and *An. gambiae* s.s. was observed after IRS withdrawal. *Anopheles arabiensis* was detected only during the IRS period. In contrast, Solenzo showed a marked increase in *An. coluzzii* proportion during and post IRS implementation, accompanied by a decline in *An. gambiae* s.s.; *An. arabiensis* was present in all periods, with a small increase after IRS. In Kampti, *An. gambiae* remained dominant across all intervention phases, while *An. coluzzii* increased after IRS implementation. *An. arabiensis* was proportionally more frequent than *An. coluzzii* during and after IRS (fig.2). Multinomial logistic regression analysis indicated that species composition varied significantly across intervention periods after accounting for site effects (Likelihood ratio test ◻^2^=21.13, df=4, p<0.001). Compared with the pre-IRS period, the odds of occurrence of *An. arabiensis* relative to *An. gambiae* s.s. were significantly higher during the IRS implementation period (OR= 4.33, 95%CI= [1.82-10.28], p= 0.0009) and remained elevated during the post-IRS period (OR= 3.87, 95%CI= [1.57-9.56], p= 0.003). In contrast, no significant temporal variation was detected for *An. coluzzii* relative to *An. Gambiae* s.s. during either the IRS implementation period (p= 0.14) or the post-IRS period (p= 0.66).

**Figure 2.**
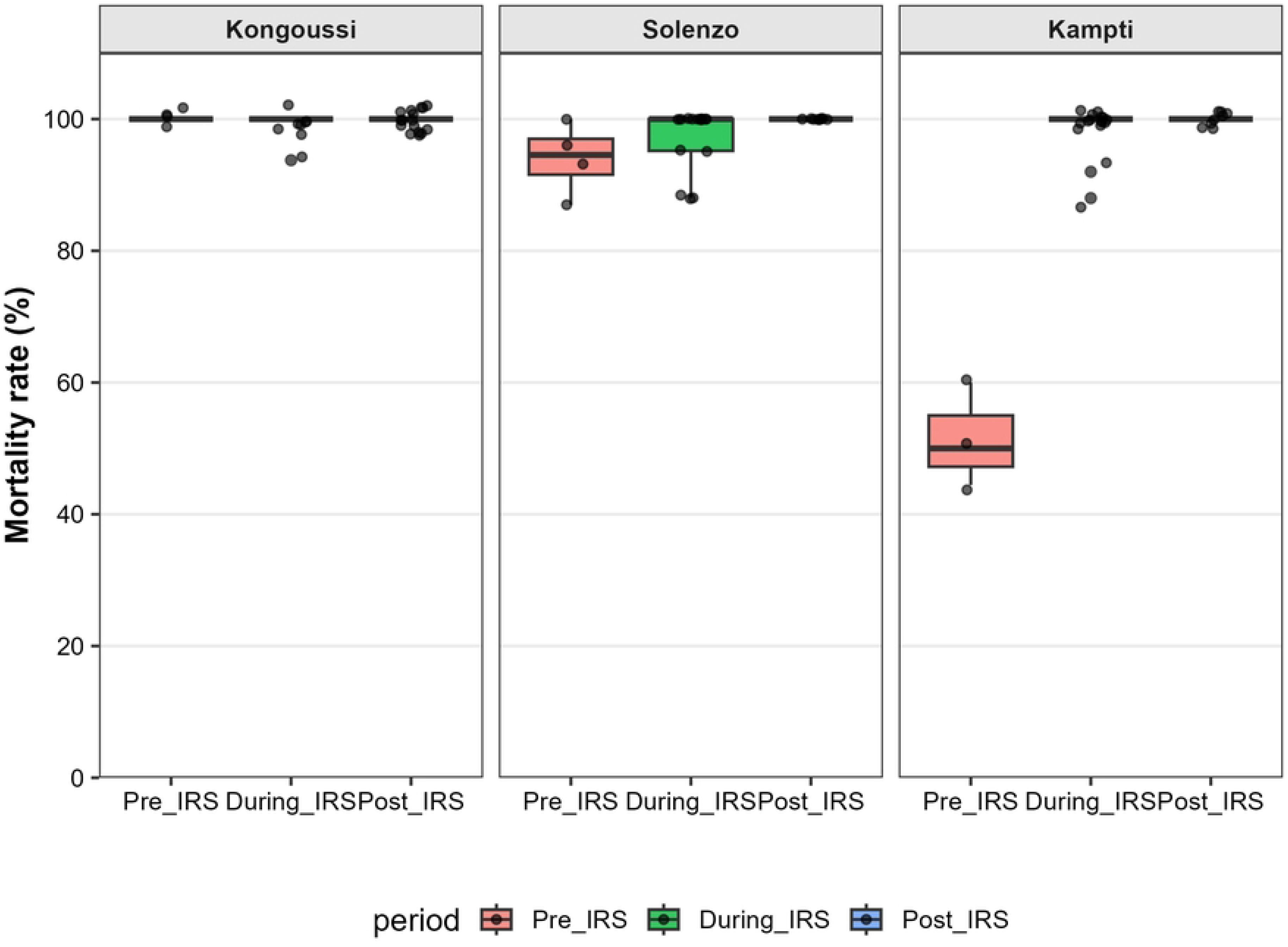
Temporal variation in species composition of the *Anopheles gambiae* complex across the three study sites. Stacked bar plots represent the relative frequencies of An. gambiae (blue), An. arabiensis (green) and An. coluzzii (orange). For each site, bars correspond respectively to the pre-IRS, during-IRS, and post-IRS periods. The sites are located in distinct ecological zones: Kongoussi (Sahelian), Solenzo (Sudan–Sahelian) and Kampti (Sudanian).

Species composition differed significantly among sites with *An. coluzzii* being substantially more frequent in Kongoussi (OR= 43.01, 95%CI: 23.93-77.33, p<0.001) and Solenzo (OR= 39.13, 95%CI: 22.23-68.89, p<0.001) than in Kampti. Furthermore, the significant interaction between site and period indicated that temporal changes in species composition differed among study sites (LRT: ◻^2^= 109.61, df= 8, p<0.001).

### Restoration of phenotypic susceptibility following IRS withdrawal

Phenotypic susceptibility assays were performed on a total of 2,015 *Anopheles gambiae* s.l. mosquitoes collected between 2017 and 2023, including 665 from Kampti (Sudanian zone), 664 from Solenzo (Sudan-Sahelian) and 686 from Kongoussi (Sahelian). The mortality rate of the *An. gambiae* s.l. populations used as controls was less than 5%, so it was not necessary to correct the mortality rates of the tested mosquitoes using Abbott’s formula. Overall, mortality increased from 86.2% before IRS period to 98.2% during IRS implementation, reaching 100% after IRS withdrawal (Fig. 3). The lower mortality observed before IRS was mainly driven by mosquito populations from Kampti. Before IRS implementation, mosquitoes from Kampti were resistant to pirimiphos-methyl with a mortality rate of 52.3%. Mortality then increased markedly to 98.8% during IRS and reached 100% after IRS withdrawal reflecting complete susceptibility according to WHO criteria. In Solenzo, mosquito populations showed suspected resistance before IRS (94% mortality) followed by high mortality during IRS (97.3%) and complete susceptibility after IRS withdrawal (100%). In contrast, mosquito populations from Kongoussi remained consistently susceptible throughout the study with mortality rate of 100% before IRS, 99.1% during IRS implementation and 100% after IRS withdrawal. Statistical analyses showed that phenotypic susceptibility varied significantly across intervention periods after accounting for site effects (LRT: χ² = 117.65, df = 2, p < 0.001). Mosquito mortality also differed significantly among study sites (LRT: χ² = 152.58, df = 6, p < 0.001). The interaction between intervention period and site was also significant (LRT: χ² = 34.93, df = 4, p < 0.001), indicating that temporal changes in phenotypic susceptibility differed between sites. Compared with the pre-IRS phase, mosquitoes collected during IRS implementation showed significantly higher odds of dying following exposure to pirimiphos-methyl (OR = 15.03, 95% CI: 8.11– 27.85, p < 0.001). No comparison could be estimated between the during-IRS and post-IRS periods because mortality reached 100% after IRS withdrawal (complete separation), which itself reflects full restoration of susceptibility according to WHO criteria. Mosquitoes collected in Kongoussi (OR = 29.92, 95% CI: 6.94–128.99, p < 0.001) and Solenzo (OR = 3.46, 95% CI: 1.83–6.56, p < 0.001) had significantly higher odds of mortality than those collected in Kampti. Collectively, these findings indicate that the restoration of susceptibility was site-dependent: the largest improvement occured in Kampti whereas populations from Kongoussi remained fully susceptible throughout the study and those from Solenzo exhibited only a modest increase in mortality.

**Figure 3.**
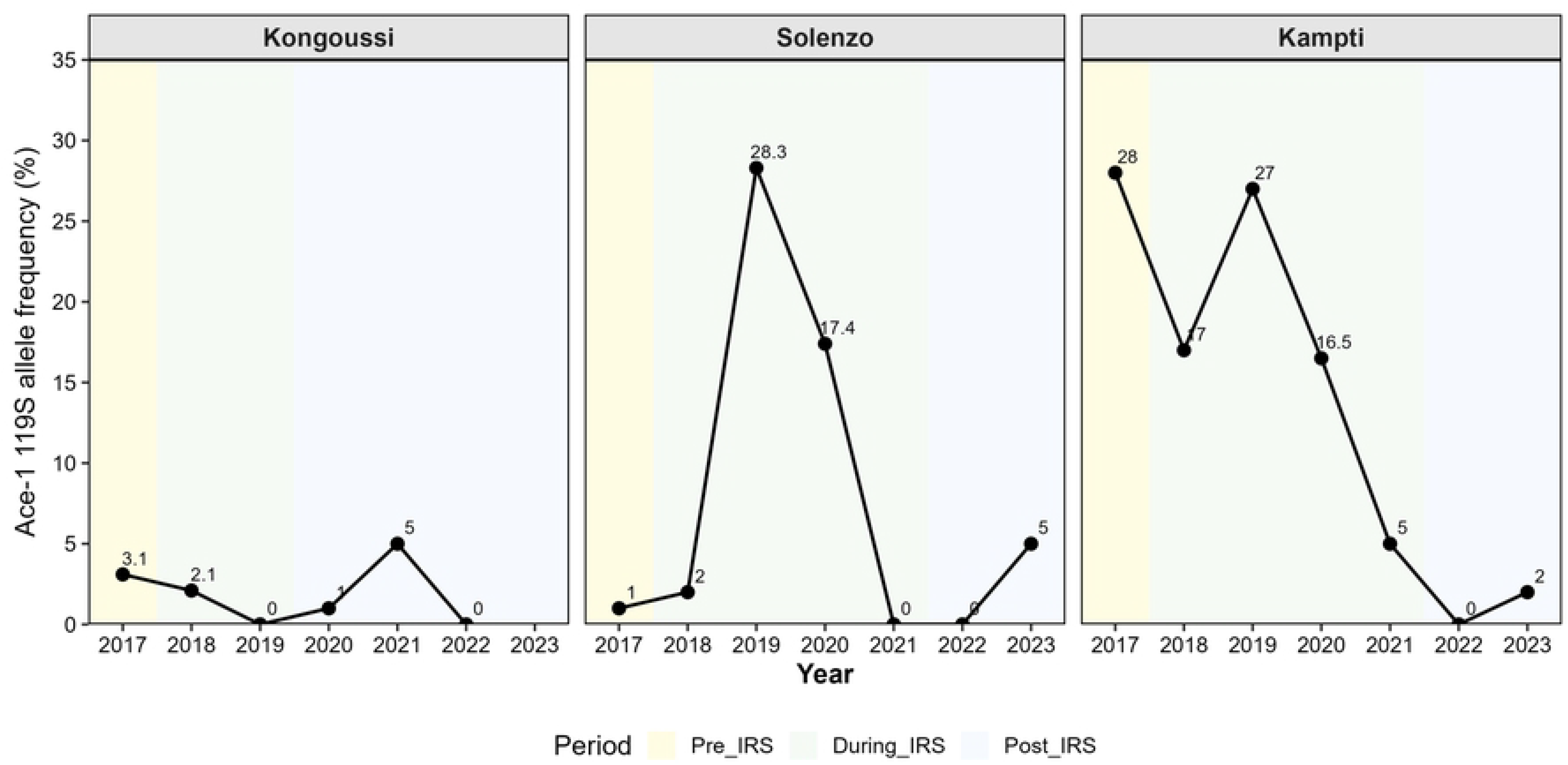
Temporal variation in *Anopheles gambiae* s.l. susceptibility across the three study sites. Boxplots represent the mortality rate observed before IRS (pink), during IRS implementation (green) and after IRS withdrawal (blue). Each point represents an individual WHO susceptibility bioassay replicate. The sites are located in distinct ecological zones: Kongoussi (Sahelian), Solenzo (Sudan–Sahelian) and Kampti (Sudanian).

### Temporal and spatial dynamics of *Ace-1* resistance allele frequencies

The same mosquito specimens that were identified to species level were subsequently genotyped for the *Ace-1* G119S mutation. Among the 1,098 specimens analyzed, 13 (1.2%) were homozygous resistant (RR), 164 (14.9%) were heterozygous (RS) and 909 (82.8%) were homozygous susceptible (SS) while 12 (1.1%) specimens yielded undetermined genotypes. The frequency of *Ace-1^R^* allele varied significantly according to year, intervention phase, study site and mosquito species (Fig. 4 and table 2). Comparison by year showed significantly higher odds of carrying the *Ace-1^R^* allele in 2019 (OR=2.31, 95%CI 1.40-3.82, p=0.001) whereas odds were significantly lower in 2021 (OR=0.33, 95%CI 0.16-0.69, p=0.003) and 2023 (OR=0.24, 95%CI 0.10-0.56, p=0.001). In 2022, no resistant allele was detected at any sites. Kongoussi maintained consistently low allele frequencies throughout the study, with only a small increase in 2021 during the post-IRS period. In contrast, Solenzo showed a pronounced increase during the IRS period, reaching its highest frequency in 2019 and 2020 before progressively declining to undetectable levels in 2021-2022 and re-emerging at low frequency in 2023 (fig.4). Kampti displayed the highest initial allele frequency in 2017 and maintained relatively elevated frequencies during the IRS period, after which the frequency declined sharply becoming undetectable in 2022 and remaining very low in 2023 after IRS withdrawal. Compared with Kampti, Kongoussi exhibited significantly lower *Ace-1^R^* frequency (OR=0.16, 95%CI 0.07-0.32, p<0.001) and moderate lower in Solenzo (OR=0.60, 95%CI 0.40-0.89, p=0.013). Species also contributed significantly to variation in *Ace-1^R^* frequency. The resistant allele occurring more frequently in *An. gambiae* than in *An. arabiensis* (OR = 2.03, 95% CI: 1.26–3.40, p = 0.005), while the increase observed in *An. coluzzii* did not reach statistical significance (p = 0.134). The interaction between year and site was significant, indicating that temporal changes in resistant allele frequency differed among sites (likelihood ratio test, p < 0.001). When comparing intervention phases, no significant difference was observed between the pre-IRS and during IRS (OR = 1.08, 95% CI: 0.72–1.68, p = 0.694). However, *Ace-1^R^* allele frequency was significantly lower during the post-IRS period compared with the pre-IRS period (OR= 0.19, 95% CI: 0.09–0.36, p<0.001) indicating a substantial reduction in the prevalence of *Ace-1* 119S allele following IRS withdrawal.

**Figure 4.**
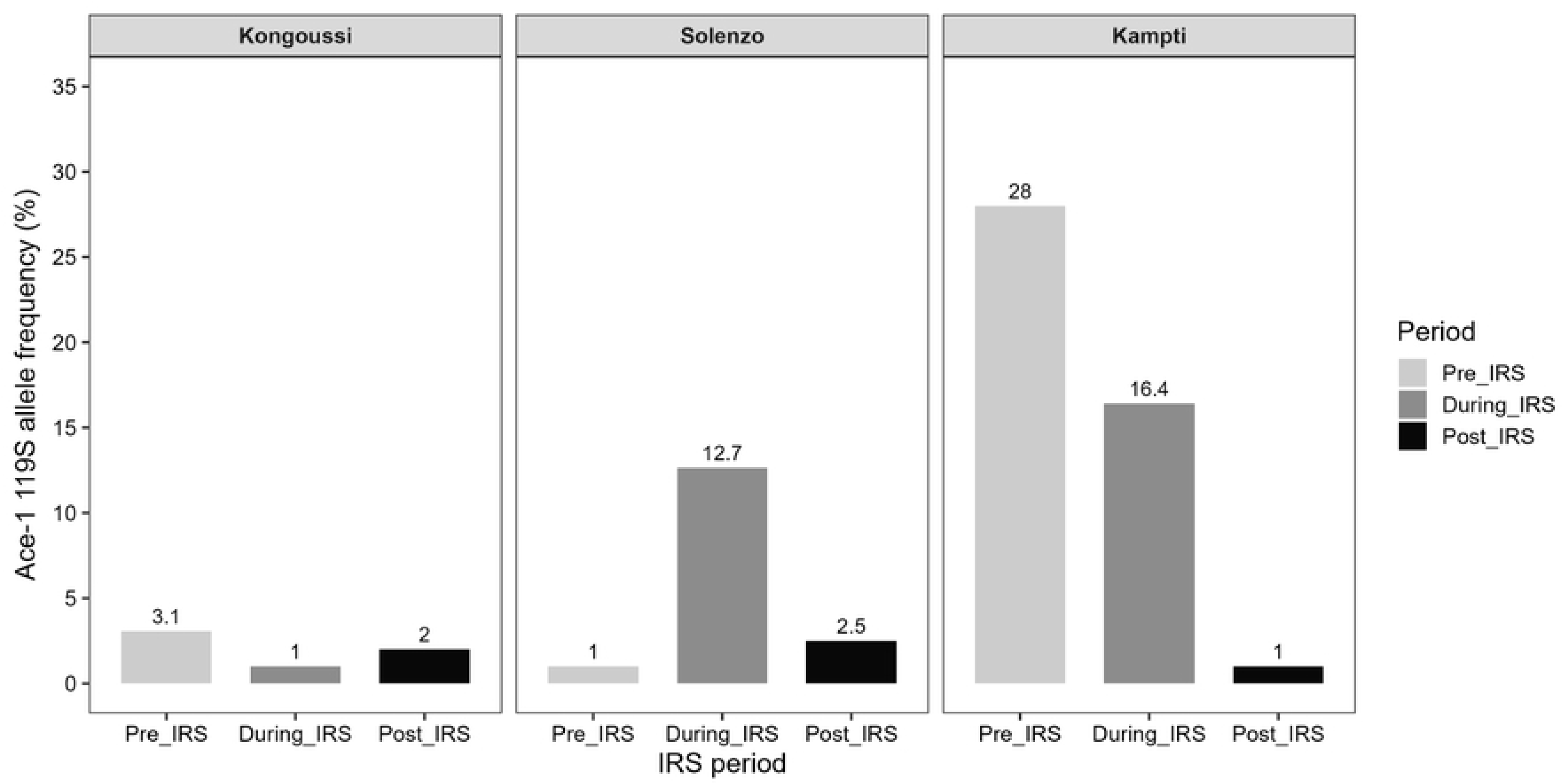
Temporal dynamic of resistant allele frequency in *Anopheles gambiae* s.l. populations across the three study sites. The frequency of the Ace-1 119S resistance allele was estimated annually for mosquito populations collected in Kongoussi, Solenzo and Kampti. Points represent the observed allele frequency. Shaded background colours correspond to the IRS implementation periods specific to each site: pre-IRS (light yellow), during IRS (light green) and post-IRS (light blue).

**Table 2.**
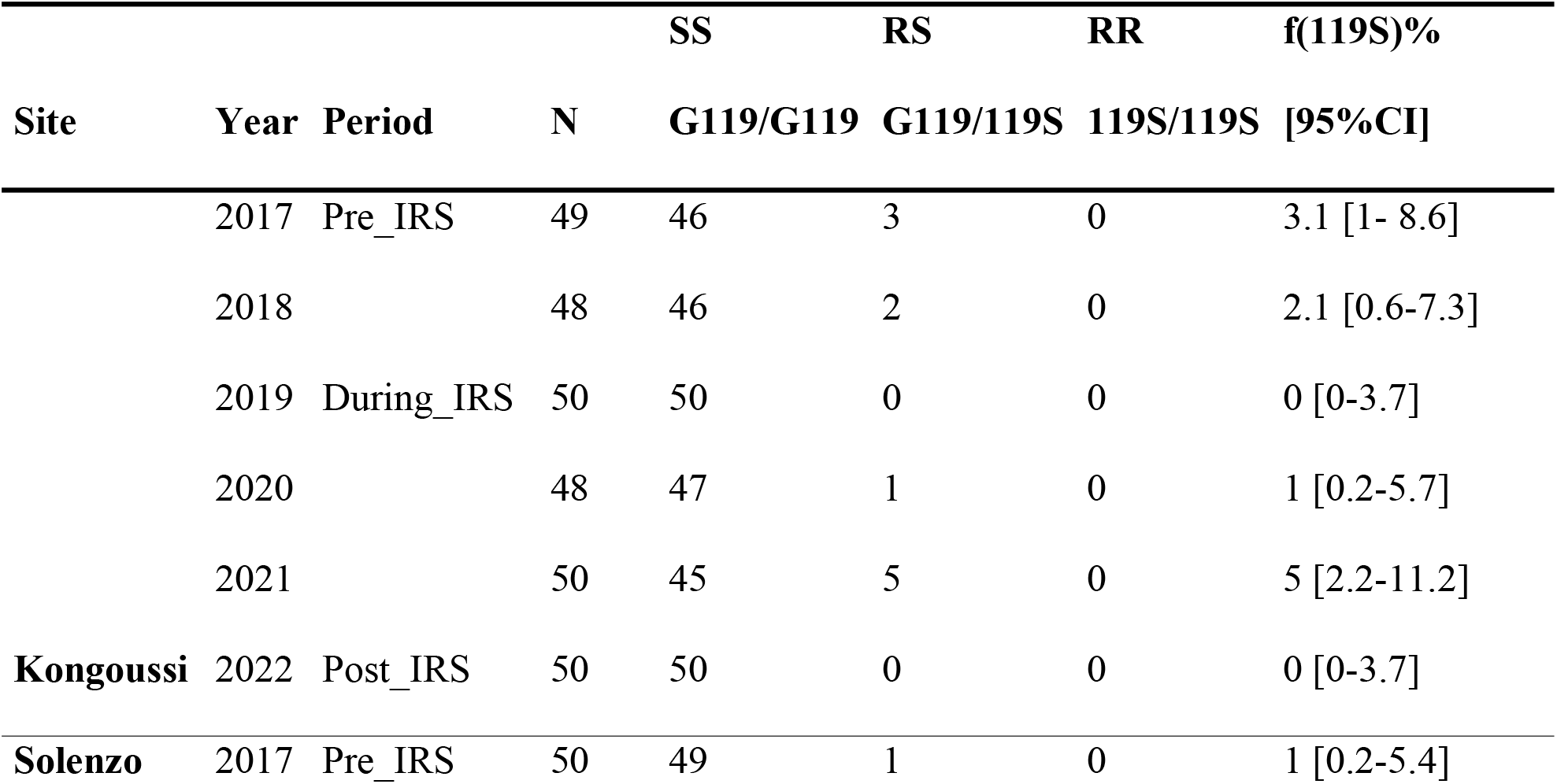
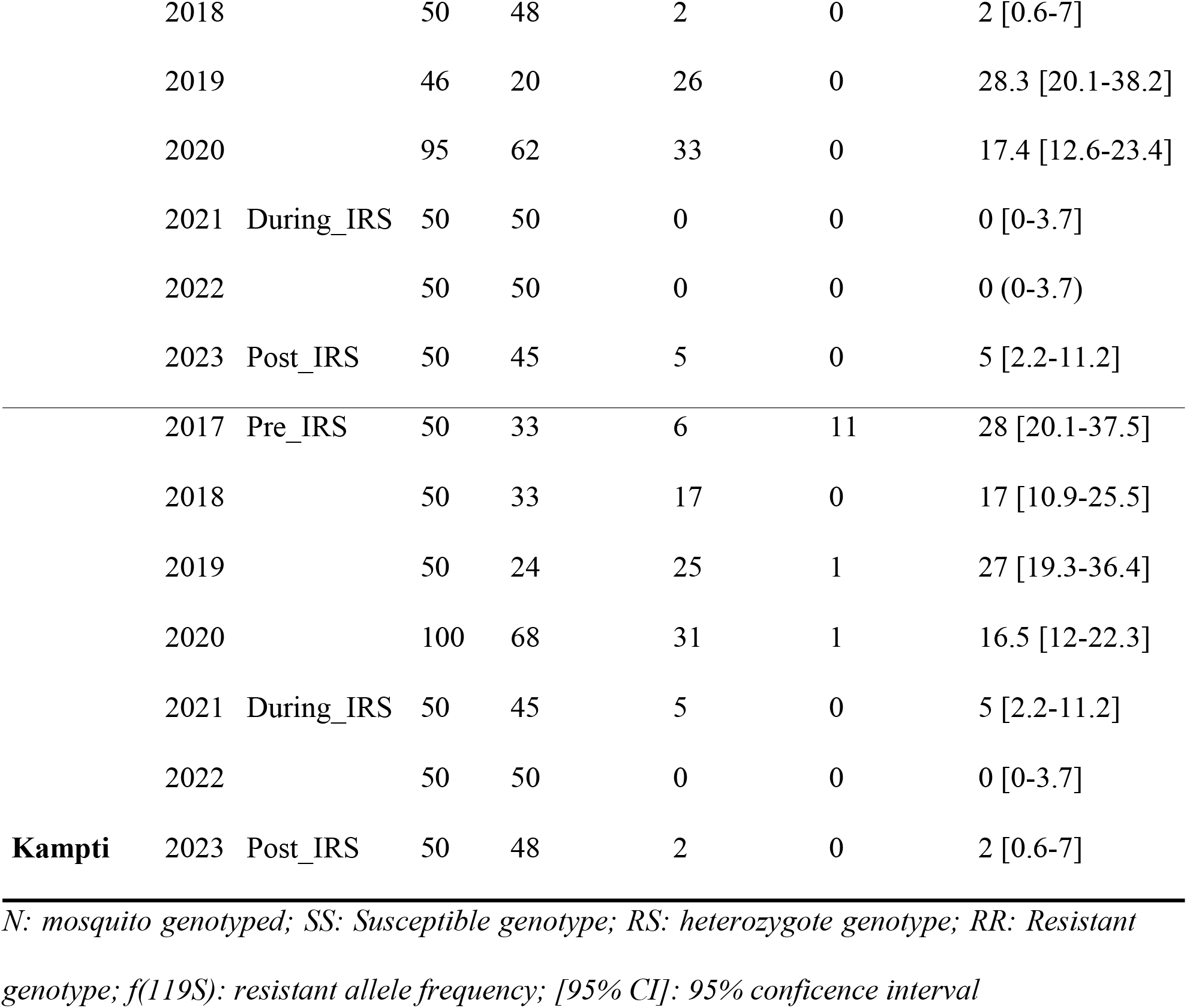
Spatial and temporal distribution of *Ace-1* G119S genotypes and resistant allele frequencies in *Anopheles gambiae* s.l. populations from study sites.

**Table 3.** Results of multivariable logistic regression models assessing the association between intervention phase, study site, mosquito species and *Ace-1* 119S allele frequency in *Anopheles gambiae* s.l. populations.

| Parameters | Variables | Odd ratio [95%CI] | p-value |
| --- | --- | --- | --- |
| **Period** | (Intercept) | 0.11 [0.06-0.20] | 2.97e-12 |
|  | Pre-IRS | Ref. | — |
|  | During-IRS | 1.08 [0.72-1.68] | 0.694 |
|  | Post-IRS | 0.19 [0.09-0.36] | 1.26e-06 |
| **Site** | Kampti | Ref. | — |
|  | Kongoussi | 0.16 [0.07-0.32] | 6.50e-07 |
|  | Solenzo | 0.60 [0.40-0.89] | 0.013 |
| **Species** | *An. arabiensis* | Ref. | — |
|  | *An. coluzzii* | 1.60 [0.87-2.99] | 0.134 |
|  | *An. gambiae* | 2.03 [1.26-3.40] | 0.005 |

*Ref: reference for comparisons*

### Decline of *Ace-1^R^* frequencies after withdrawal of IRS

The frequencies of the *Ace-1* 119S resistant allele varied by study site and by the periods during which IRS was implemented (Figure 5). A marked decrease in the frequency of the resistant allele was observed after the discontinuation of IRS in Kampti, where it fell from 28% during the pre-IRS period to just 1% during the post-IRS period. A similar trend, although less pronounced, was observed in Solenzo where the frequencies of the *Ace-1* 119S allele increased during the IRS implementation period before decreasing markedly after IRS withdrawal. In contrast, Kongoussi showed consistently low frequencies of the resistant allele throughout the study period, with only minor fluctuations between the different intervention periods (Figure 5).

**Figure 5.**
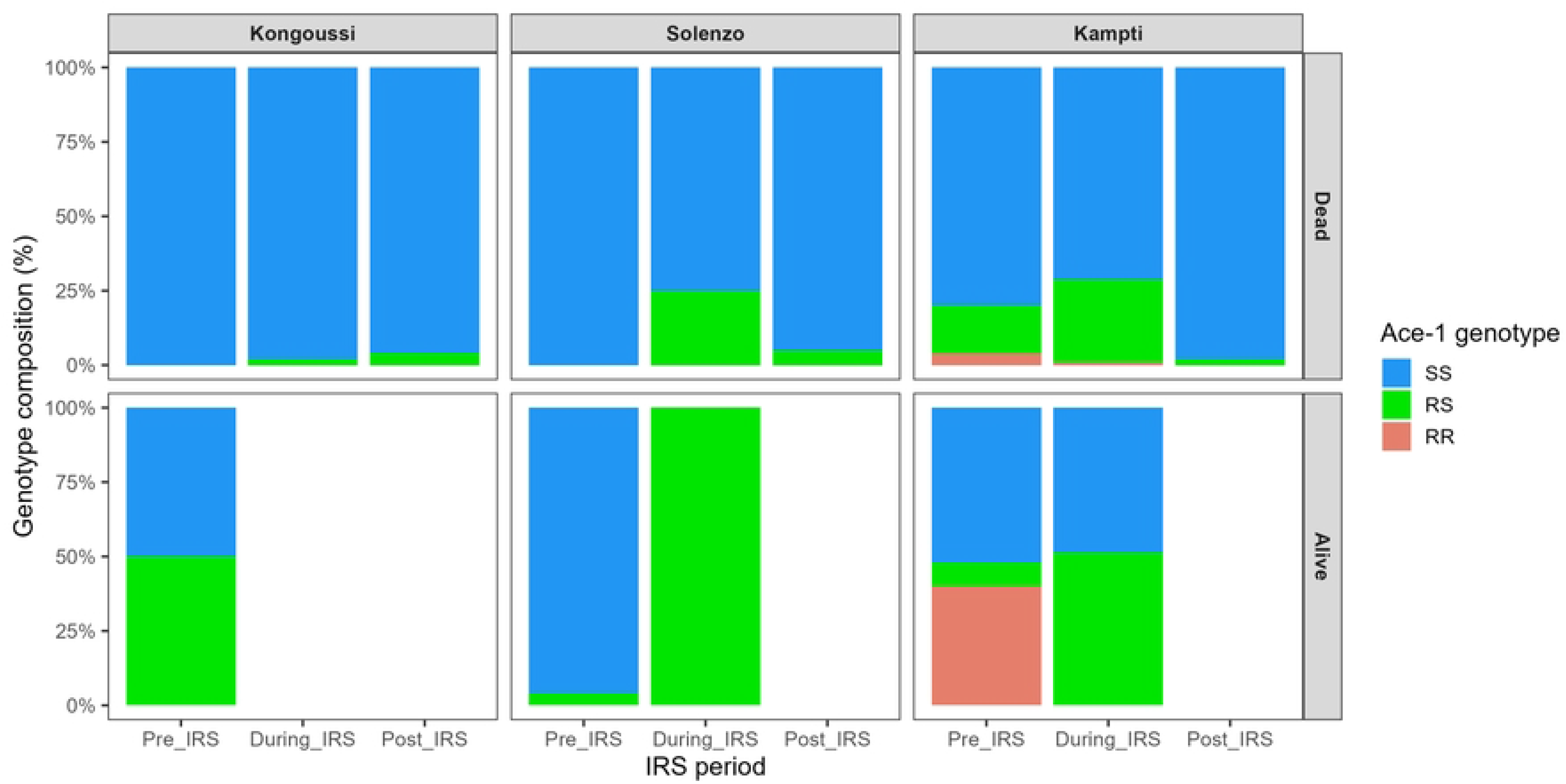
Variation of *Ace-1* 119S allele frequency across IRS implementation periods in *Anopheles gambiae* s.l. populations from study sites. The frequency of the Ace-1 119S resistance allele was estimated by period for mosquito populations collected in Kongoussi, Solenzo and Kampti. Bars represent the observed allele frequency at each period: pre-IRS (dark gray), during IRS (medium gray) and post-IRS (light gray).

### Genotype–phenotype relationships

The distribution of *Ace-1* genotypes differed markedly between mosquitoes that died and those that survived following exposure to pirimiphos-methyl (Figure 6). Across all study sites and IRS periods, mosquitoes that died after exposure were predominantly homozygous susceptible (SS). Heterozygous resistant mosquitoes (RS) represented a smaller proportion of dead individuals, whereas homozygous resistant mosquitoes (RR) were rare and were detected exclusively in Kampti. In contrast, among mosquitoes that survived exposure, resistant genotypes were proportionally more frequent. Surviving mosquitoes were predominantly heterozygous resistant (RS), while homozygous resistant individuals (RR) constituted a substantial proportion of survivors in Kampti, particularly during the pre-IRS period. Following IRS withdrawal, no mosquitoes survived exposure to pirimiphos-methyl at any of the study sites and homozygous resistant individuals were no longer detected among tested populations. Statistical analyses revealed a strong association between *Ace-1* genotype and survival following exposure to pirimiphos-methyl. Compared with homozygous susceptible mosquitoes (SS), heterozygous resistant mosquitoes (RS) exhibited significantly higher odds of surviving exposure (OR = 1.83, 95% CI: 1.09–3.08, p = 0.023). The survival advantage was considerably stronger for homozygous resistant mosquitoes (RR), which were approximately 28 times more likely to survive exposure than SS individuals (OR = 27.62, 95% CI: 6.91–110.45, p < 0.001). Mosquito species also influenced survival probability. Compared with *An. gambiae*, both *An. arabiensis* (OR = 0.24, 95% CI: 0.10–0.58, p = 0.002) and *An. coluzzii* (OR = 0.46, 95% CI: 0.23–0.91, p = 0.025) showed significantly lower odds of surviving exposure to pirimiphos-methyl. Similarly, mosquitoes collected in Kongoussi exhibited significantly lower survival probabilities than those from Kampti (OR = 0.20, 95% CI: 0.08–0.49, p < 0.001), whereas survival probabilities in Solenzo did not differ significantly from those observed in Kampti (p = 0.168).

**Figure 6.** Distribution of *Ace-1* genotype among dead and surviving mosquitoes according to IRS implementation period and study site. The frequency of the Ace-1 genotype was estimated by period for mosquito populations collected in Kongoussi, Solenzo and Kampti. Barplots represent at each period the observed genotype: homozygote susceptible (light blue), heterozygote resistant (light green) and homozygote resistant (light red).

### Concordance between molecular and phenotypic resistance indicators

A significant negative correlation was observed between *Ace-1* 119S allele frequency and mortality following exposure to pirimiphos-methyl (Spearman’s ρ= −0.48, p = 0.033), indicating that mosquito populations with higher frequencies of the resistant allele tended to exhibit lower mortality rates. This concordance between molecular and phenotypic indicators supports the role of *Ace-1*-mediated resistance in determining susceptibility to pirimiphos-methyl.

### Evidence supporting fitness costs associated with *Ace-1*-mediated resistance

Several converging observations obtained during the longitudinal study provide evidence supporting the existence of biological costs associated with *Ace-1*-mediated resistance in natural populations of *An. gambiae* s.l. in Burkina Faso. First, a gradual restoration of phenotypic susceptibility to pirimiphos-methyl was observed after the cessation of IRS campaigns. Mortality rates recorded during WHO bioassays gradually increased over time, reaching 100% at all sites once the IRS campaigns had ended, indicating an apparent disappearance of phenotypic resistance to organophosphates. Second, this restoration of susceptibility was accompanied by a marked decrease in the frequency of the *Ace-1* 119S resistance allele after the cessation of interventions, particularly at the Kampti and Solenzo sites. In Kampti, the mean allele frequency dropped from 28% before IRS to about 1% after IRS, while in Solenzo, the high frequencies observed during the implementation of the IRS decreased sharply after the removal of the selection pressure exerted by pirimiphos-methyl. Third, resistant genotypes, particularly resistant homozygotes (RR), became rare or undetectable once IRS campaigns were discontinued. Fourth, genotype-phenotype analyses showed that mosquitoes carrying the resistant allele had a significant survival advantage after exposure to pirimiphos-methyl, with homozygous resistant (RR) individuals having a survival probability approximately 28 times higher than that of homozygous susceptible (SS) individuals, while heterozygotes (RS) also exhibited a significantly higher survival probability. Fifth, a significant negative correlation was observed between the frequency of the *Ace-1* 119S allele and mortality rates recorded in pirimiphos-methyl susceptibility bioassays (Spearman’s correlation: ρ = −0.48; p = 0.033). Populations with low frequencies of the resistant allele were thus associated with higher mortality rates following exposure to the insecticide. Taken together, these results are consistent with the hypothesis that the reduction in selection pressure exerted by indoor spraying has contributed to the decline of resistant genotypes in the natural populations studied.

## DISCUSSION

### Main findings

This longitudinal study evaluated whether the changes observed after the discontinuation of IRS corresponded to the biological costs associated with the *Ace-1* resistance allele. Overall, the results showed a gradual restoration of phenotypic susceptibility to pirimiphos-methyl following the discontinuation of the interventions, accompanied by a significant decrease in the frequency of the *Ace-1* 119S resistance allele and a decline in the prevalence of resistant genotypes, particularly resistant homozygotes (RR). Despite the survival advantage conferred by the resistant allele under exposure to pirimiphos-methyl, its frequency declined sharply after the cessation of IRS campaigns, suggesting a loss of its selective advantage in the absence of insecticide pressure. Our results also highlight significant spatial heterogeneity in resistance dynamics among the study sites. While Kampti and Solenzo exhibited relatively high frequencies of the *Ace-1* 119S allele during the years when IRS was implemented, Kongoussi maintained low frequencies throughout the study period. These differences suggest that local resistance dynamics likely result from complex interactions between the intensity of selection pressure, the specific composition of vector populations and local ecological contexts.

### Withdrawal of IRS and restoration of susceptibility

The gradual recovery of susceptibility to pirimiphos-methyl after the cessation of indoor residual spraying campaigns is one of the main findings of this study. *Anopheles* populations that were resistant or suspected resistant during the first years of follow-up gradually regained high mortality rates, reaching full susceptibility according to WHO criteria during the post-IRS period. This trend is consistent with the theoretical principles of insecticide resistance management, according to which the removal or reduction of selection pressure can promote the restoration of susceptibility when resistance mechanisms no longer confer an adaptive advantage in the given environment [1, 9]. Similar trends in the restoration of susceptibility following the withdrawal of insecticides have been reported in several species of insect vectors and pests when resistance was based primarily on mechanisms involving modifications of the molecular target or certain metabolic mechanisms associated with biological costs for the carriers [8, 9, 10]. In the specific context of Burkina Faso, this recovery may also have been facilitated by the rotation of insecticides used in national vector control strategies, which potentially reduced the use of organophosphates for vector control [17]. Furthermore, local changes in agricultural practices or in the non-public-health use of insecticides may also have contributed to reducing the selection pressures exerted on mosquito populations. Taken alone, however, the restoration of susceptibility observed here does not demonstrate a biological cost associated with *Ace-1*-mediated resistance. What makes that interpretation plausible is the concordance between phenotypic trends, changes in allele frequencies, and the gradual disappearance of resistant genotypes following the withdrawal of interventions.

### Declining *Ace-1* frequencies indicate relaxation of selection pressure

In addition to restoring phenotypic susceptibility, our study revealed a significant decrease in the frequency of the *Ace-1* 119S resistant allele following the cessation of indoor spraying campaigns, particularly at the Kampti and Solenzo sites. This decrease was accompanied by a gradual decline in the abundance of resistant genotypes, notably resistant homozygotes (RR), which were virtually absent during the post-IRS period. These results are consistent with the predictions of evolutionary theory, which predict that the frequencies of resistance alleles selected under insecticide pressure may decrease when that pressure is removed or greatly reduced, provided that they no longer confer a selective advantage in the given environment [9]. In the case of organophosphates and carbamates, the *Ace-1* G119S mutation alters the target site of acetylcholinesterase, allowing resistant mosquitoes to survive insecticide exposure but potentially also impairing the enzyme’s normal physiological function. The sharp decline in frequencies observed in Kampti and, to a lesser extent, in Solenzo thus suggests that, following the withdrawal of Actellic® 300SCS, susceptible genotypes have gradually regained a competitive advantage within natural populations. In contrast, Kongoussi maintained low frequencies throughout the study, likely due to historically lower selection pressure resulting from a shorter duration of IRS implementation at this site, and possibly to more limited agricultural use of organophosphates in this area, partly as a consequence of insecurity. Similar patterns of declining *Ace-1* frequencies following a reduction in selection pressures have already been reported in several regions of West Africa [9, 18]. However, the sporadic persistence of low residual frequencies observed at some sites in our study suggests that other factors, such as agricultural use of insecticides, mosquito population movements, or genetic introgression, could contribute to the maintenance of the resistant allele at low levels within local populations [18, 21].

### Evidence of fitness costs associated with *Ace-1* resistance

The central hypothesis of this study was that discontinuing indoor spraying with pirimiphos-methyl would lead to a decrease in *Ace-1* frequencies if resistance mediated by this mutation were associated with biological costs for its carriers. Several observations made during this study are consistent with this hypothesis. First, genotype-phenotype analyses confirmed that mosquitoes carrying the resistant allele had a significant survival advantage when exposed to pirimiphos-methyl. Resistant heterozygotes (RS) had a significantly higher probability of survival than susceptible homozygotes, while resistant homozygotes (RR) had the highest survival probabilities. Despite this advantage under insecticide pressure, the frequencies of resistant genotypes declined sharply after the discontinuation of IRS campaigns. Furthermore, the gradual disappearance of resistant homozygotes after the cessation of treatments is consistent with the existence of selective disadvantages associated with these genotypes in the absence of exposure to organophosphates. In *Culex pipiens,* insecticide resistance genes have been shown to improve a mating competitiveness cost on resistant males, which favors the return of susceptible genotypes once insecticide treatments are discontinued [19].

In *An. gambiae*, several experimental and field studies have also suggested the existence of biological costs associated with the *Ace-1* G119S mutation. In particular, studies have shown that duplications of the *Ace-1* locus can mitigate some of these costs by allowing the coexistence of a susceptible copy and a resistant copy of the gene, thereby reducing the deleterious effects associated with the resistant mutation alone [6, 10]. Similarly, the work of Grau-Bové et al. [3] showed that the spread of resistance to organophosphates in West Africa was largely driven by duplication and introgression of the *Ace-1* locus among species of the *An. gambiae* complex. Biological costs associated with resistance mechanisms have also been reported in *An. funestus*, particularly for metabolic resistance mediated by glutathione S-transferases and cytochrome P450 enzymes, although the underlying mechanisms differ from those involving the modification of the target site observed for *Ace-1* [8]. We did not measure the classical components of fitness-fecundity, fertility, longevity or mating competitiveness-directly, but inferred them from the evolutionary trajectories observed in natural populations following the removal of the selective pressure exerted by pirimiphos-methyl. Consequently, our results provide field evidence consistent with the existence of biological costs associated with *Ace-1*-mediated resistance, rather than a direct experimental demonstration of these costs.

### Why did resistance not disappear completely?

Despite the sharp decline in the frequency of the *Ace-1* 119S allele observed after the cessation of indoor spraying, resistance has not completely disappeared from the populations studied. Low residual frequencies of the resistant allele persisted at certain sites, notably in Kampti and Solenzo, suggesting that additional mechanisms likely contribute to the maintenance of resistance in the absence of direct pressure from public health interventions. One possible explanation lies in the use of insecticides from the same chemical class in agriculture. In Burkina Faso, several studies have highlighted significant pesticide use in cotton, vegetable, and cereal crops, which may expose larval and adult *Anopheles* populations to various insecticides and thus maintain residual selection pressure on resistance mechanisms [20]. Although the organophosphates used in agriculture are not necessarily identical to the pirimiphos-methyl used in public health, the phenomena of cross-resistance or co-selection cannot be ruled out.

The movement of mosquito populations between geographic areas could also contribute to the maintenance of low residual frequencies of *Ace-1^R^* in local populations. Gene flow between populations subjected to different selection pressures can regularly reintroduce resistant alleles into areas where selection has become weak or absent, thereby slowing their complete disappearance. Recent genomic studies conducted in Burkina Faso have also shown that insecticide-resistance variants continue to spread within *An. gambiae* s.l. populations through evolutionary processes involving gene flow, recombination, and the emergence of new haplotypes associated with resistance [21]. These processes may contribute to the persistence of resistant alleles even after the selection pressure exerted by insecticides has diminished. Finally, duplication of the *Ace-1* locus is likely one of the most plausible explanations for the residual persistence of resistance. Studies conducted in West Africa have shown that heterogeneous duplications of the *Ace-1* locus allow for the coexistence of a susceptible copy and a resistant copy of the gene, thereby reducing the biological costs associated with the G119S mutation while retaining part of the selective advantage under insecticide pressure [3, 6, 10]. In the absence of information on duplications at this locus in our study, their potential contribution to the persistence of the low frequencies observed cannot be ruled out.

### Implications for insecticide resistance management

Our results provide important insights for strategies to manage insecticide resistance in Burkina Faso and, more broadly, in sub-Saharan Africa. The rapid decline in *Ace-1* frequencies after spraying was discontinued suggests that rotating or alternating insecticides may help partially restore the susceptibility of vector populations when resistance mechanisms are associated with significant biological costs. These observations support international recommendations for integrated resistance management strategies based on the rotation of insecticide families and the combined use of multiple vector control tools [1, 12]. In this context, alternating between indoor residual spraying using different classes of insecticides and the deployment of new-generation insecticide-treated bed nets could help limit the long-term accumulation of resistance alleles within natural populations. The gradual replacement of conventional pyrethroid-based mosquito nets with nets containing PBO, followed by combinations with chlorfenapyr, combined with a rotation of the insecticides used in indoor spraying, could thus constitute a particularly promising approach for sustainably preserving the effectiveness of available vector control tools.

### Study limitations

This study has several limitations that must be taken into account when interpreting the results. First, the biological costs associated with *Ace-1*-mediated resistance were not measured directly but were inferred from the evolutionary trajectories observed in natural populations after the removal of the selective pressure exerted by pirimiphos-methyl.

Second, no direct measurements of the classical components of fitness such as fecundity, fertility, longevity or mating competitiveness were made in resistant mosquitoes. Such data would have allowed for a more precise characterization of the nature and magnitude of the costs associated with the different resistance genotypes. Third, our study did not include analyses to identify the possible presence of duplications at the *Ace-1* locus, now recognized as a major mechanism capable of mitigating the biological costs associated with the G119S mutation. Fourth, the annual sample sized available for genotyping (approximately 50 mosquitoes per site and year) were modest, o the allele-frequency estimates for individual years carry wide confidence intervals. Finally, other environmental factors that could shape the observed dynamics agricultural use of insecticides, mosquito population movements and local variation in ecological conditions were not quantified.

## CONCLUSIONS

This longitudinal study of natural populations of *An. gambiae* s.l. in Burkina Faso shows that, following the cessation of indoor residual spraying campaigns, the frequency of the *Ace-1* 119S resistance allele declined sharply, particularly at sites where resistance levels were highest during the intervention period. This trend was accompanied by a gradual restoration of phenotypic susceptibility to pirimiphos-methyl, with mosquito populations returning to mortality levels consistent with complete susceptibility according to WHO criteria. The correlation between the decline in allele frequencies, the gradual disappearance of resistant genotypes, and the restoration of phenotypic susceptibility provides field evidence consistent with the existence of biological costs associated with *Ace-1*-mediated resistance in *An. Gambiae* s.l. These findings highlight the importance of taking evolutionary processes into account in the management of insecticide resistance. They provide valuable insights to guide future resistance management strategies in Burkina Faso and other regions where malaria is endemic and where organophosphate-based interventions may be implemented.

## Acknowledgments

We would like to express our gratitude to all the staff at the Institute for Health Sciences Research (IRSS)/Western Regional Directorate (DRO) for their support during larval collection, molecular analyses, and other joint activities. We also extend our sincere thanks to the partners of the PMI VectorLink project, as well as to the staff of Burkina Faso’s health districts, for their collaboration and support, which contributed to the completion of this study.

## Availability of data and materials

Data are fully available from the corresponding author upon request.

## Author Contributions

**Conceptualization:** RKD

**Data curation:** RKD

**Formal analysis**: DOK ASH DDS

**Funding acquisition**: RKD

**Investigation**: RKD

**Methodology**: RKD ASH DDS DOK DAK LKN SM BGM RB

**Project administration:** RKD

**Resources**: RKD

**Software**: DDS ASH

**Validation**: RKD ASH DDS

**Visualization**: DOK ASH DDS LKN DPAK SM RB BGM NM RKD

**Writing - original draft**: DOK ASH DDS RKD

**Writing - review & editing**: DOK ASH DDS RKD

